# The effects of emotional valence and intensity on cognitive and affective empathy after insula lesions

**DOI:** 10.1101/2021.03.28.436842

**Authors:** Olga Holtmann, Marcel Franz, Constanze Moenig, Jan-Gerd Tenberge, Christoph Preul, Wolfram Schwindt, Maximilian Bruchmann, Nico Melzer, Wolfgang H.R. Miltner, Thomas Straube

## Abstract

The insula plays a central role in empathy. However, the complex structure of empathic deficits following insular damage is not fully understood. While previous lesion research has shown variable deficits in patients with insular damage on basic discrimination tasks or self-report measures, it is unclear in how far patients with insular damage are impaired in cognitive (CE) and affective empathy (AE) functions depending on valence and arousal of stimuli using an ecologically valid paradigm. In the present study, patients with insular lesions (*n* = 20) and demographically-matched healthy controls (*n* = 24) viewed 16 videos (duration: 60 sec each) that varied in terms of valence and emotional intensity. The videos showed a person (target) reporting on a personal life event. In CE conditions, subjects continuously rated the affective state of the target, while in AE conditions they continuously rated their own affect. Mean Squared Error (MSE) assessed deviations between subject and target ratings (CE: deviation between targets’ and participants’ ratings of targets’ emotions; AE: deviation between targets’ and participants’ self-ratings of emotion). Patients differed from controls only in negative, low intensity AE, rating their own affective state less negative than the target rated his/her affect. This deficit was not related to trait empathy, neuropsychological or clinical parameters, or laterality of lesion. Our findings provide important insights into the profile of social cognition impairment after insular damage. Empathic functions may be widely spared after insular damage in a naturalistic, dynamic setting, potentially due to the intact interpretation of social context cues by residual networks outside the lesion. The particular role of the insula in AE for negative states may evolve specifically in situations that bear higher uncertainty, which points to a threshold role of the insula in online ratings of AE.

## 1. Introduction

Empathy is fundamental for social interaction (Hall et al., 2016) and lack of empathy is a distinguishing mark in a number of psychopathologies (Hillis, 2014; Kumfor et al., 2017; Shamay-Tsoory, 2011). Empathy is a complex multidimensional construct including affective and cognitive components (Zaki & Ochsner, 2012). Cognitive empathy (CE), or perspective taking, refers to the correct inference of emotional states in others (Shamay-Tsoory & Lamm, 2018; Zaki & Ochsner, 2012). Affective empathy (AE), or experience sharing, is the ability to vicariously respond with the same emotion to the emotional state of another (Shamay-Tsoory & Lamm, 2018; Zaki & Ochsner, 2012). Distinct, though potentially overlapping, networks have been suggested in AE and CE (Hillis, 2014; Shamay-Tsoory & Lamm, 2018; Zaki & Ochsner, 2012). Yet, the role of specific brain regions in AE and/or CE is not fully understood.

The insula has a role in empathy (Decety, 2015; Engen & Singer, 2013; Gu et al., 2013; Hillis, 2014). A substantial body of functional neuroimaging research points to a particular involvement in AE (Banissy et al., 2012; Marsh, 2018; Shamay-Tsoory, 2011; Shamay-Tsoory & Lamm, 2018; Singer et al., 2009; Wicker et al., 2003; Zaki & Ochsner, 2012), due to the role of the insula in interoception, generation of emotional experience and emotional awareness (Gu et al., 2013; Menon & Uddin, 2010; Shamay-Tsoory, 2011; Uddin et al., 2017; Walter, 2012; Zaki & Ochsner, 2012). The insula is consistently activated in vicarious responses in the domain of disgust, pleasant or unpleasant odours or tastes, physical and emotional pain, fear, anxiety, happiness, and other social emotions such as embarrassment or admiration (for reviews and meta-analyses see Bernhardt & Singer, 2012; Gogolla, 2017; Gu et al., 2013; Kurth et al., 2010; Uddin et al., 2017). In a meta-analysis of fMRI studies on empathy, Fan et al. (2011) found the right insula associated with AE, and the left insula with AE and CE. This finding is in line with previous evidence suggesting the insula as a neural interface for self-awareness, social cognition and the sensorimotor system (Adolfi et al., 2017; Uddin et al., 2017). Functional connectivity analyses found that CE is associated with functional connectivity between the anterior insula, superior temporal sulcus, medial prefrontal cortex and the brainstem (Gallagher & Frith, 2003). However, based on fMRI findings alone, the causal role of the insula in empathy functions cannot be solved (Hillis, 2014; Kumfor et al., 2017).

Studies of patients with focal insula damage offer a unique opportunity to specify whether the insula is required in AE and/or CE. While lesion-based research has already explored some emotional and socio-cognitive constructs that may affect empathic functions (e.g., emotion experience, emotion expression, facial emotion recognition) (Adolphs et al., 2003; Borg et al., 2013; Boucher et al., 2015; Calder et al., 2000; Terasawa et al., 2015), only a handful of studies have addressed effects of insular damage on empathy tasks. The findings are mixed. Some studies support the critical role of the insula in AE: Driscoll et al.(Driscoll et al., 2012) found that self-reported AE correlated with insular lesion volume in a sample of patients with penetrating brain injury. Gu et al.(2012) found deficits in implicit and explicit pain perception in three patients with anterior insular damage. In contrast, Leigh et al. (2013) found deficits of emotional inference associated with infarcts in the anterior insula in a sample of patients with acute right-hemisphere ischemic stroke. Others suggest a general role of the insula in empathy: Couto et al. (2013) reported reduced emotion evaluation of spontaneous facial expressions and reduced comprehension of pain situations (besides a multimodal emotion-recognition deficit) in a patient with focal right-hemispheric damage of white-matter association tracts adjacent to the insula. Chen et al. (2016) found multidimensional empathy deficits (self-report AE and CE, facial emotion recognition, pain perception, emotion inference in social interaction scenes) in patients with insula gliomas, as compared to healthy and lesion controls. However, these studies need to be interpreted with caution since they (1) have been based on small samples with heterogeneous insula damage patterns, (2) have not systematically investigated AE and CE dysfunctions in one paradigm, (3) have used a specific operationalization of AE in terms of affect sharing (Coll et al., 2017), (4) have used discrimination tasks with static, staged stimuli and discrete response options or self-report measures that may not fully represent actual empathic abilities because of limited ecological validity (Coll et al., 2017; Kumfor et al., 2017; Zaki & Ochsner, 2009). There is evidence of poor correlations between ecologically valid measures of empathy and discrimination tasks or self-report measures (Harvey et al., 2013; Kumfor et al., 2017). Notably, some studies of neuropsychiatric samples suffering from insular dysfunction (e.g. frontotemporal lobar degeneration, or Huntington’s disease) reported emotion recognition impairments on forced-choice recognition tests, which were absent when patients were tested with more ecological tasks providing contextual cues (for review see Kumfor et al., 2017)).

Psychological frameworks conceptualize emotion along two dimensions: valence (level of experience pleasantness/ unpleasantness) and arousal (level of intensity of a given emotion) (Lang, 2014; Posner et al., 2005). As with emotional experience, empathy can be differentiated according to valence and arousal (Morelli et al., 2015). The insula has a role in valence and arousal processing (Craig, 2002; Critchley, 2005; Kurth et al., 2010). Previous lesion studies have documented impaired evaluation of emotional valence and arousal (Berntson et al., 2011), apathy and emotional blunting after insular damage (Ibañez et al., 2010; Jones et al., 2010). Processing of negative emotions may be particularly affected by insular damage (Calder et al., 2000; Couto et al., 2013; Holtmann et al., 2020; Terasawa et al., 2015; Vicario et al., 2017). Although previous lesion studies of empathy support the deficit bias towards negative emotions (Couto et al., 2013; Gu et al., 2012a), systematic investigations of empathic processing according to valence and arousal are lacking. Thus, it is unclear in how far patients with insular damage are impaired in CE and AE functions depending on valence and emotional intensity of stimuli in an ecologically valid paradigm.

In the present study, we examined patients with unilateral insular damage (*n*=20) as well as healthy controls (*n*=24) on a modified empathy accuracy task (Zaki et al., 2009; Zaki & Ochsner, 2009) that included naturalistic stimuli and, thus, accounted for the dynamic nature of empathy. For the paradigm, we created and validated a new video set consisting of 16 video clips (duration: 60 sec each) in German language that varied in terms of valence (positive vs. negative) and emotional intensity (high intensity vs. low intensity). The videos showed a person (‘target’) speaking of a personal life event. During each film clip, subjects were instructed to continuously rate changes in emotional valence of either the target’s emotion or his/her own emotion. To outline response biases in patients, we computed the Mean Squared Error (MSE) to quantify deviations between subject and target ratings. For CE trials, we compared targets’ and subjects’ ratings of targets’ emotions. For AE trials, we compared targets’ and subjects’ self-ratings of emotions, thus employing a new behavioral index of AE (Coll et al., 2017; Morrison et al., 2016). Based on previous literature, we assumed to find disproportionate changes in the patient group relative to the control group for AE, particularly for negative stimuli. If deficits represent a threshold effect, that is patients need a higher intensity for normal empathic responses, then the differences between controls and patients should be more pronounced for low intense stimuli.

## 2. Materials and methods

### 2.1. Participants

Twenty patients (6 females) with a history of either left (*n* = 13) or right (*n* = 7) damage centred on the insula were enrolled in the study. Patients were recruited to the research programme through the Department of Neurology, University Hospital Muenster, Muenster, Germany (*n* = 11), or following participation in a rehabilitative programme (Miltner et al., 1999, 2016) run by the Department of Clinical Psychology, Friedrich Schiller University Jena, Jena, Germany, and the Department of Neurology, University Hospital Jena, Jena, Germany (*n* = 9). Inclusion criteria were as follows: (1) unilateral lesions due to middle-cerebral-artery stroke, centrally affecting the insula, as confirmed by three experts in clinical neuroimaging (CP, WS and NM) from CT or MRI scans of the brain; (2) stable lesions (at least 1 year after lesion onset); (3) no cognitive deficits compromising the understanding of task instructions and task performance (i.e., global aphasia, attention deficits, amnesia, disorders of reasoning, or visual neglect); (4) no history of neurodegenerative disorders, epilepsy, brain tumours, or brain trauma; (5) no history of substance-induced disorders; (6) no history of psychiatric disorders; (7) motor ability to participate in the experimental procedure. Single case and group-level demographical and clinical data are presented in Table 1. The left and the right-lesioned group did not differ in terms of sex, education, age at testing, or lesion age (all *P* ≥ 0.110). On average, total lesion volume (*P* = 0.870) and anatomical distribution of damaged tissue were similar between the patient groups (see Fig.1 for the overlap of reconstructed lesions, Supplementary Fig. 1 for individual anatomical images, and Supplementary Table 1 for detailed lesion analysis). Comprehensive neuropsychological assessment targeting major neurocognitive domains was conducted (see Table 1). Left and right-injured patients did not differ on neuropsychological characteristics (all *P* ≥ 0.138). Visual field examination provided no evidence for manifest neglect (i.e. no blind spots).

**Table 1:** Single-case and group-level patient characteristics.

| Patient | Clinical |  |  | Demographics |  |  | Performance on neuropsychological tests (z-scores) |  |  |  |  |  |  |  |  | Depression |
| --- | --- | --- | --- | --- | --- | --- | --- | --- | --- | --- | --- | --- | --- | --- | --- | --- |
|  | Lesion | Lesion | Lesion | Age | Sex | Education | Verbal IQ | Nonverbal IQ | Visuo-spatial | Auditory Memory | Auditory Memory | Phasic | Selective | Divided | Visual field/Neglect | BDI-II |
|  | laterality | age | volume | (years) |  | (years) | (WAIS-IV-Sim) | (WAIS-IV-MR) | memory | (immediate recall) | (delayed recall) | alertness | attention | attention | (TAP-VFCT, blind spots) |  |
|  |  | (years) | (cm<sup>3</sup>) |  |  |  |  |  | (CFT-CQM) | (WMS-IV-LMi) | (WMS-IV-LMd) | (TAP-Alert) | (TAP-Go/NoGo) | (TAP-DA) |  |  |
| L1 | left | 8 | 2.4 | 66 | M | 13 | -1.00 | 1.67 | 0.80 | -0.67 | -0.67 | -1.70 | 0.20 | 1.30 | 0 | 12 |
| L2 | left | 6 | 13.4 | 54 | M | 13 | -2.33<sup>a</sup> | 0.00 | 1.60 | -1.67 | -0.67 | 0.50 | 0.30 | 1.00 | 0 | 15 |
| L3 | left | 12 | 64.8 | 51 | M | 13 | -0.67 | 0.67 | 0.10 | -0.33 | -0.33 | -1.70 | 0.20 | 0.90 | 0 | 2 |
| L4 | left | 3 | 14.5 | 44 | M | 12 | 0.00 | 1.00 | 0.30 | 0.67 | 1.67 | -0.92 | 0.31 | 0.81 | 0 | 1 |
| L5 | left | 6 | 24.8 | 53 | M | 9 | -2.33<sup>a</sup> | -1.67 | 1.60 | -1.00 | -1.33 | 0.10 | 0.31 | -0.10 | 0 | 17 |
| L6 | left | 1 | 8.9 | 55 | M | 12 | 0.67 | -1.00 | 0.50 | 0.67 | 1.00 | 0.00 | -1.17 | -1.28 | 0 | 23<sup>b</sup> |
| L7 | left | 4 | 30.6 | 56 | M | 10 | 1.00 | 0.67 | 0.30 | -1.33 | -0.67 | 0.20 | 0.31 | -1.64 | 0 | 24<sup>b</sup> |
| L8 | left | 1 | 1.5 | 56 | M | 9 | 0.00 | 0.67 | 0.50 | -2.33<sup>a</sup> | -1.67 | -2.05<sup>a</sup> | 0.20 | -0.10 | 0 | 11 |
| L9 | left | 8 | 34.6 | 64 | M | 13 | -0.33 | 0.33 | -1.50 | 0.33 | 0.67 | -1.28 | -1.48 | -0.61 | 0 | 6 |
| L10 | left | 1 | 1.3 | 73 | M | 10 | 1.00 | 0.33 | -0.60 | -1.33 | -0.33 | -- | -- | -- | 0 | 8 |
| L11 | left | 4 | 33.2 | 52 | F | 13 | -1.67 | -1.33 | -0.90 | -2.33<sup>a</sup> | -2.33<sup>a</sup> | -1.90 | 0.30 | -1.70 | 0 | 15 |
| L12 | left | 10 | 18.7 | 65 | F | 13 | 0.00 | -1.00 | -1.20 | -0.33 | 0.00 | 2.30 | 0.30 | 0.00 | 0 | 22<sup>b</sup> |
| L13 | left | 4 | 9.1 | 58 | F | 13 | 0.00 | 2.00 | -1.10 | -0.33 | -1.00 | -0.20 | 0.30 | -0.10 | 0 | 9 |
| R1 | right | 6 | 29.1 | 67 | M | 13 | 1.33 | 0.33 | 0.10 | -0.33 | -1.00 | -0.20 | -0.90 | -0.60 | 0 | 4 |
| R2 | right | 6 | 19.7 | 57 | M | 9 | -0.67 | -0.33 | 0.30 | 1.00 | 0.33 | -1.75 | 0.20 | -1.28 | 0 | 6 |
| R3 | right | 18 | 31.2 | 69 | M | 9 | -1.33 | 0.00 | -0.10 | 2.00 | 0.00 | 0.31 | 0.20 | -0.61 | 0 | 3 |
| R4 | right | 1 | 11.1 | 71 | M | 9 | 0.67 | 1.00 | 0.60 | -1.67 | -0.33 | -1.41 | -0.41 | 1.75 | 0 | 6 |
| R5 | right | 4 | 7.3 | 55 | F | 10 | -1.00 | -1.67 | -1.30 | -1.00 | -1.67 | 0.00 | 0.20 | -1.50 | 0 | 12 |
| R6 | right | 8 | 15.2 | 60 | F | 12 | 1.00 | 1.00 | 0.10 | -0.67 | -1.00 | 0.10 | 0.20 | 1.20 | 0 | 27<sup>b</sup> |
| R7 | right | 1 | 33.8 | 63 | F | 13 | 0.67 | 1.33 | 0.70 | 1.00 | 2.33 | -1.75 | 0.20 | 1.17 | 0 | 12 |
| **Left-lesioned patients** |  | 5.2 ± 3.5 | 19.8 ± 17.9 | 57.5 ± 7.7 | 10M/ 3F | 11.8 ± 0.5 | -0.44 ± 1.13 | 0.18 ± 1.14 | 0.03 ± 1.02 | -0.77 ± 1.01 | -0.44 ± 1.09 | -0.55 ± 1.28 | 0.01 ± 0.63 | -0.13 ± 1.03 | 0 | 12.7 ± 7.6 |
| **Right-lesioned patients** |  | 6.3 ± 5.8 | 21.1 ± 10.4 | 63.1 ± 6.1 | 4M/ 3F | 10.7 ± 0.7 | 0.10 ± 1.07 | 0.24 ± 1.03 | 0.06 ± 0.66 | 0.05 ± 1.31 | -0.19 ± 1.30 | -0.67 ± 0.92 | -0.04 ± 0.44 | 0.02 ± 1.32 | 0 | 10.00 ± 8.3 |
| **P** |  | 0.616 | 0.870 | 0.110 | 0.613 | 0.209 | 0.319 | 0.911 | 0.952 | 0.138 | 0.659 | 0.835 | 0.853 | 0.792 | -- | 0.472 |

*Note:* Group-level data are presented as means and standard deviations for each parameter, except for absolute frequencies provided for sex. BDI-II = Beck Depression Inventory, CFT-CQM = memory quotient from the Rey Complex Figure Test, TAP = Test Battery for Attention Performance, TAP-Alert = TAP-subtest 'Alertness', TAP-Go/NoGo = TAP-subtest 'Go/NoGo' (number of errors), TAP-DA = TAP-subtest 'Divided attention I, aud-vis' (number of errors), TAP-VFCT = TAP-subtest 'Visual field with central task', WAIS-IV-Sim = subtest 'Similarities' from the Wechsler Adult Intelligence Scale (4<sup>th</sup> edition), WAIS-IV-MR = subtest 'Matrix Reasoning' from the Wechsler Adult Intelligence Scale (4<sup>th</sup> edition), WMS-IV-LMi = subtest 'Logical Memory', immediate recall, from the Wechsler Memory Scale (4<sup>th</sup> edition), WMS-IV-LMd = subtest 'Logical Memory', delayed recall, from the Wechsler Memory Scale (4<sup>th</sup> edition), SD = standard deviation, F = female, M = male
Statistical comparisons: t-tests, Fisher's exact probability test (sex)
<sup>a</sup> 2 SD below the mean of normative controls as provided by individual test manuals, <sup>b</sup> above the clinical cut-off of 18 for clinical depression

**Figure 1:**
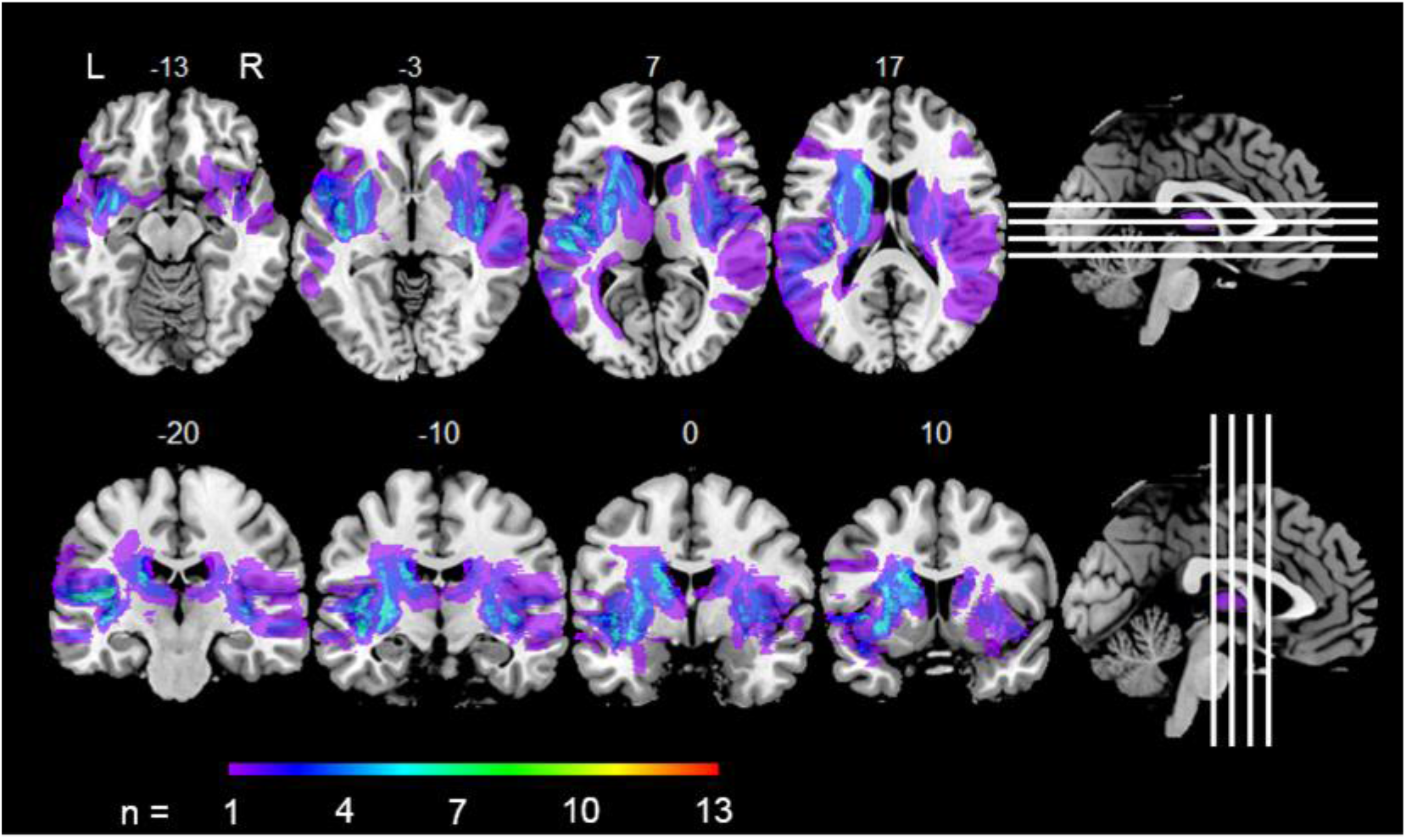
Overlay map of individual lesions determined in both patient groups, superimposed onto a T_1_-weighted MNI template 1×1×1 mm^3^, with MNI coordinates of each axial (z-axis, upper panel) and coronal section (y-axis, lower panel) provided for lesion localization. Colour bar shows the number of patients with a lesion in the respective voxel (L = left hemisphere, overlap among left-lesioned patients; R = right hemisphere, overlap among right-lesioned patients).

Twenty-four (12 females) neurologically and psychiatrically healthy controls were recruited via public announcements in Jena (*n* = 12) and Muenster (*n* = 12). There were no statistical differences between patients and controls on demographical characteristics (age: [*M* ± *SD*], 59.5 ± 7.5 years [patients] vs. 58.6 ± 7.7 years [controls], *P* = 0.723; level of education: 11.4 ± 1.8 years [patients] vs. 11.9 ± 1.4 years [controls], *P* = 0.296; sex: *P* = 0.227). Controls had significantly lower scores on the Beck-Depression Inventory-II (BDI-II(Beck et al., 1996)), as compared to patients (11.8 ± 7.7 [patients] vs. 7.3 *±* 5.8 [controls], *P* = 0.034). All subjects had normal or corrected-to-normal visual acuity.

Informed written consent was obtained from each participant, and financial compensation was given for participation. The experimental procedures conformed to the Declaration of Helsinki and were approved by and performed in accordance with the guidelines of the Ethics Committee of the German Society of Psychology (TS032013_rev). All participants were right-handed.

### 2.2. Lesion reconstruction

High-resolution isotropic (1 mm) 3D structural T_1_-weighted imaging data were obtained in 14 patients using a 3 Tesla system (Magnetom Prisma, Siemens, Erlangen, Germany) within 40 days of participation in the study. Six patients did not provide consent for MRI or did not meet the inclusion criteria for MRI at the time of testing. In these cases, T_2_FLAIR-weighted MR images (3mm) obtained at 1.5 Tesla (Philips Achieva, Philips Medical Systems, Best, The Netherlands) or cranial CT images (5 mm; Somatom Definition Flash Siemens Medical Solutions, Forchheim, Germany; H30s soft kernel) from the latest clinical routine follow-up screening were used for lesion analyses. The anatomical images were aligned to Montreal Neurological Institute (MNI) space (1 × 1 × 1 mm^3^) by means of the Clinical Toolbox for SPM12 (Wellcome Trust, London, UK), which includes a specialized standard MRI and CT template and spatial normalization algorithms that are recommended for studies of stroke-aged populations (Rorden et al., 2007; Rorden & Brett, 2000; Rorden et al., 2012). Individual lesion boundaries were identified by means of the Clusterize Toolbox (version 1.0beta) available for SPM12, offering a semi-automatic approach to lesion demarcation regardless of image modality (Clas et al., 2012; de Haan et al., 2015). During the automated preprocessing procedure, each image was inspected for local intensity maxima, thus defining cluster cores. Minimum size for the initiation of a cluster was set to 150 mm^3^ and a lower intensity threshold of 20% was used for the elimination of irrelevant background voxels (de Haan et al., 2015). The intensity threshold was then iteratively adapted in steps of 1%, resulting in the assignment of each voxel to a cluster core. Clusters corresponding to damaged tissue were then interactively selected and, if necessary, modified by changing the intensity threshold. Finally, resulting lesion demarcations were reviewed and manually adjusted by a professional neurologist (CM) who was blind to the study’s hypotheses, data and statistical analyses at this time. MriCron (http://www.mccauslandcenter.sc.edu/crnl/tools) was used to create within-group overlaps and displays on a T_1_-weighted MNI template (Rorden et al., 2007). Regional extent of damaged tissue was determined by using standard anatomical atlases implemented in MriCron, i.e. Automated Anatomic Labelling (AAL) atlas (Tzourio-Mazoyer et al., 2002) and JHU white matter tractography atlas (Oishi et al., 2008).

### 2.3. Empathy task

#### 2.3.1 Video acquisition and selection

Twenty-nine native German-speaking adult volunteers (19 females, mean age = 25.4 years, range: 20-44 years) participated as targets in exchange for monetary compensation. All of them provided a written informed consent prior to participation. Targets were asked to recall at least four autobiographic events, in which they experienced either a positive or a negative emotion in either high or low affective intensity. For each suggested event they were willing to discuss while being filmed, targets wrote a short summary and provided ratings of affective valence (9-point scale, from −4, ‘very negative’ to 4, ‘very positive’), overall affective intensity (9-point scale, from 1, ‘no emotion’ to 9, ‘very strong emotion’) and experienced intensity for several provided emotional categories, with the opportunity to add own categories (9-point scale, from 1, ‘not felt at all’ to 9, ‘felt most intensely’). Emotional category ranked first for a particular event was considered as target emotion. The experimenter used these ratings to compile a set of events differing in valence, intensity and content to be filmed. Targets were then seated in front of a video camera that was positioned slightly behind the experimenter to approximate a naturalistic interpersonal interaction. The camera captured a frontal image of the target’s face and shoulders against a white background, which allowed a high resolution of facial signals. Targets received a list with the events put in a randomized order for filming. Before starting the recording of each event, targets read the respective summary and were given time to recreate sensory and affective experiences of their memory (Mackes et al., 2018; Zaki et al., 2009). When they were ready, the filming started. Each clip lasted for approximately 60 sec. After each filming, targets repeated ratings of valence, intensity and specific emotional experience for the affective state during their discussion of the event (Zaki et al., 2009). Upon completion of the recording procedure, targets watched each of their videos and this time continuously rated their affective valence in the video while using the same 9-point scale as above. Starting point for each continuous rating was ‘0’, i.e. neutral position. This continuous valence rating served as the “true” emotional response to which perceivers’ ratings were later compared. Finally, targets were asked for their consent to use their videos as part of the empathic accuracy task, and all of them provided a written permission.

Out of 203 recorded videos, we selected 43 videos that matched a series of criteria: (i) content appropriateness for different age groups, (II) representation of diverse emotional categories, and (III) discriminative ability regarding affective valence (positive vs. negative) and affective intensity (high vs. low) of content as determined from target ratings obtained after each recording. The 43 video vignettes were piloted with a small sample of healthy participants (*n* =10, 5 females, mean age = 25.4, range: 20-35 years). Based on target and pilot ratings, we selected two final sets á eight video vignettes to be used in the present study. Detailed description of characteristics is provided in Supplementary Table 2. Each set was divided equally between female and male targets, positive and negative valence, as well as high-intensity and low-intensity contents. As determined from target and pilot ratings, both sets were capable to discriminate valence and intensity, with no statistical differences between the two sets observed (for details see Supplementary Table 3). For the following experimental procedure, one set was assigned to conditions of affective empathy, while the other set was used only in conditions of cognitive empathy. Two videos of different targets than in the main experiment were selected for pre-training purposes.

#### 2.3.2. Procedure

Participants completed a modified version of the empathic accuracy task (Zaki et al., 2008, 2009). In line with the original empathic accuracy protocol (Zaki et al., 2008, 2009), participants were instructed to continuously rate the perceived affective valence of the target, using the same 9-point scale as targets themselves had used (i.e., from −4, ‘very negative’ to 4, ‘very positive’). This rating procedure is used to infer cognitive empathic accuracy (Brook & Kosson, 2013; Coll et al., 2017). Extending the original protocol, we asked the participants to continuously judge their own affect in response to target videos on the same valence scale as above. This rating procedure has been suggested to tap affective empathic accuracy (Coll et al., 2017; Morrison et al., 2016). In both types of trials, participants were asked to focus on “moment-to-moment-changes” in the affective state of the target/ self. Default starting point for each rating was ‘0’, i.e. neutral position. Videos were presented in four blocks (two for cognitive empathy, two for affective empathy), thus aiming to minimize potential influences between the different rating procedures (Coll et al., 2017). Presentation of blocks was randomized, with no repetition of blocks belonging to the same rating procedure. Four videos (one positive video of high affective intensity, one positive video of low affective intensity, one negative video of high affective intensity, and one negative video of low affective intensity) were assigned to each block; they appeared only within this block, but in a randomized order. In each video trial, a cue was presented for 5 sec, followed by the video and concomitant rating, followed by a fixation cross presented for 15 sec. Cues were rating instructions preparing the participant for the upcoming rating procedure. In conditions of affective empathy, the cue consisted of the sentence “How do you feel while hearing this narrative?”; in conditions of cognitive empathy the cue was “How does the person in the video feel?”. Before starting the main experiment, participants were carefully trained in both rating procedures. They watched and rated one practice video for each rating procedure, and the experimenter verified that the task was understood properly. Completion of the empathic accuracy task required approximately 30 min. Stimulus presentation and response recording were controlled by Presentation software Version 19.0 (Neurobehavioral Systems, Inc.).

### 2.4. Self-reported empathy

Patients and controls completed the German adaptation of the Interpersonal Reactivity Index (IRI) (Davis, 1983), the Saarbrücker Persönlichkeitsfragebogen (SPF; Paulus, 2009), after the empathic accuracy task. This widely used self-report questionnaire addresses individual differences in cognitive and affective trait empathy by means of four subscales: The Perspective-Taking (PT) scale and the Fantasy (FS) scale represent the concept of cognitive empathy, while the Empathic-Concern (EC) scale and the Personal-Distress (PD) scale operationalize affective empathy. The mean score of the PT and the FS subscales was used to quantify cognitive empathy, while the mean score of the EC and PD subscales was employed as a measure of affective empathy (Chen et al., 2016).

### 2.5. Statistical procedures

The aim of the present work was to explore response biases exhibited by patients with insular damage on cognitive resp. affective empathy tasks, as a function of valence and affective intensity. To this end, we compared subject with target ratings by using the Mean Squared Error (MSE). MSE scores were obtained for each subject and for each video. For each video, the subject and the target rating were averaged across one-second intervals, so that each one-second mean served as one point in the time-series analysis. MSE scores were computed by subtracting the target from the subject rating for each point in the time series, squaring the difference values and averaging the squared difference values across the time series. MSE scores from cognitive resp. affective empathy trials were highly related to coefficients of empathic accuracy (*r* = −0.85, *P* < 0.001) resp. empathic congruence (*r* = −0.82, *P* < 0.001), as derived from correlating target and subject rating profiles according to previous research employing the Empathic Accuracy Task (Morrison et al., 2016; Zaki et al., 2008, 2008, 2009). However, while correlation coefficients emphasize solely the type and extent of mutual fluctuation of target and subject ratings, MSE scores add information on the absolute deviation between target and subject rating profiles, which aids the quantification of response biases. MSE scores were subjected to a 2 (Group: patients, controls) × 2 (Valence: positive, negative) × 2 (Intensity: high, low) mixed-design analysis of covariance (ANCOVA), with Group as between-subjects factor, Valence and Intensity as within-subject factors, and BDI-II as the covariate, since patients and controls significantly differed on the BDI-II (see Participants Section). Bonferroni-Holm-corrected *post-hoc* tests were used to investigate the nature of significant interactions. To extend the insight into significant response biases in the patient group, we analysed whether, on average, target profiles were under- or overestimated by the subjects. For each subject and for each video in question, we integrated the area under the curve of difference values (rating subject – rating target) by application of the trapezoidal rule. Positive Area Under the Curve (AUC) scores showed that the subject responded more positive than the target (rating subject > rating target). Negative AUC scores indicated that the subject reacted more negative than the target (rating subject < rating target). AUC scores were subjected to a one-way ANCOVA with Group as between-subject factor, and BDI-II as the covariate. Data reduction and computation of MSE and AUC were performed using Matlab 9.1.0 (The MathWorks Inc., Natick, Massachussetts, United States).

SPF scores were examined separately for the cognitive and affective empathy subscale, using a 2 (Group) × 2 (Subscale: PT and FS for cognitive empathy, or EC and PD for affective empathy) mixed-design ANCOVA, with Group as between-subject factor, and Subscale as within-subject factor. No *post-hoc* testing was conducted, since no significant interactions occurred.

Two-sided, independent *t*-tests were used to compare left and right-lesioned patients on MSE scores or total scores for cognitive resp. affective empathy from the SPF. Pearson’s product-moment correlations were used to explore relationships between MSE scores and self-report data, separately in patients and controls. Pearson’s product-moment correlations were also used to explore whether significantly altered MSE scores in patients were related to cognitive measures. Statistical significance was assumed when *P* < 0.05. Statistical analyses were performed using IBM SPSS Statistics (Version 26, IBM Corp., Armonk, New York, United States).

### 2.6. Data availability

The datasets generated during and/or analysed during the present study are partially included in this article (Supplementary Material) or are available from the corresponding author on reasonable request.

## 3. Results

### 3.1. Empathy task

Cognitive empathy was indexed by the MSE between targets’ and subjects’ ratings of the targets’ affective state. A mixed-design ANCOVA revealed a significant main effect of Valence (F(1,41) = 6.33, *P* = 0.016, *η^2^* = 0.13), indicating significantly larger deviations between target and subject ratings for positive than for negative stimuli. We found no significant main effect of Intensity (F(1,41) = 0.87, *P* = 0.356), or Group (F(1,41) = 0.18, *P* = 0.676). The Group by Valence interaction (F(1,41) = 0.01, *P* = 0.946), the Group by Intensity interaction (F(1,41) = 0.96, *P* = 0.334), and the Group by Valence by Intensity interaction (F(1,41) = 2.35, *P* = 0.133) were not significant. Thus, compared to healthy controls, patients with insular lesions did not reveal any difficulties to continuously track emotions in targets. Mean MSE scores from cognitive empathy trials are shown in Fig. 2A, as a function of group (patients or controls), valence (positive or negative) and affective intensity (high or low) of the stimuli.

**Figure 2:**
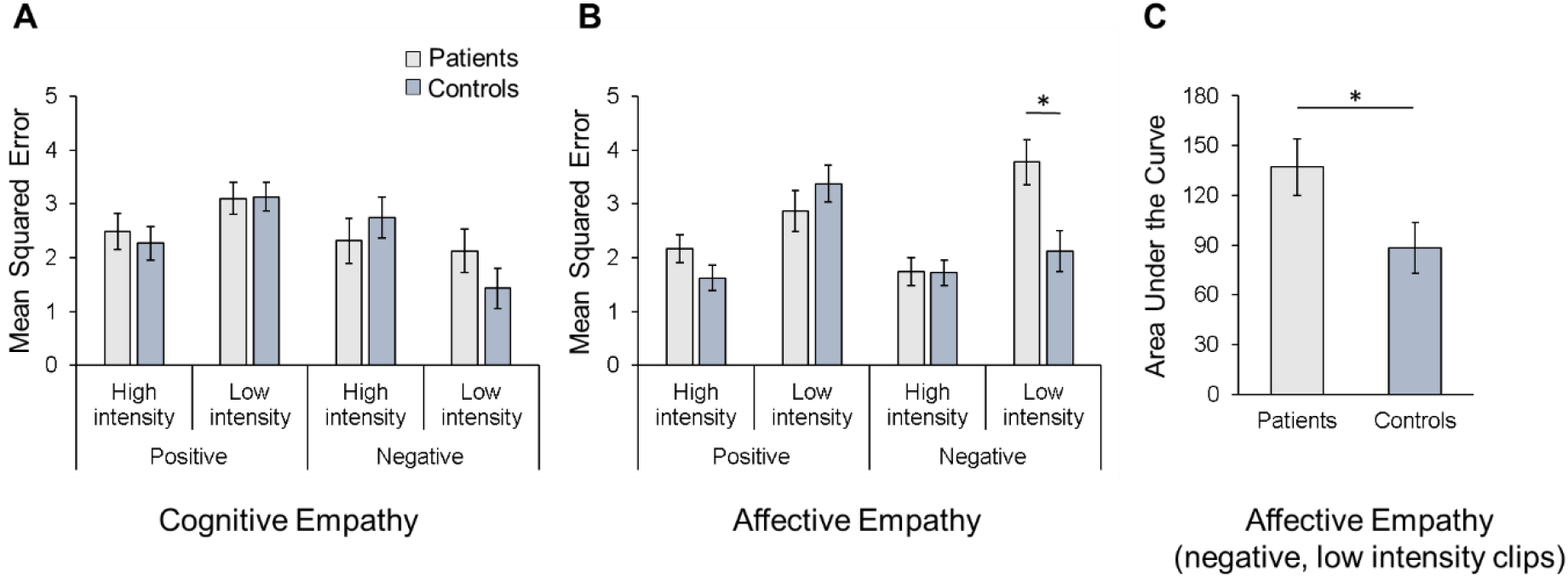
Mean squared error (MSE) for cognitive empathy trials(**A**) and affective empathy trials (**B**) in patients with insular lesions and healthy controls by valence (positive or negative) and affective intensity (high or low). Higher MSE scores represent larger absolute deviations between target and subject ratings (cognitive empathy: ratings of targets’ affect; affective empathy: self-ratings of affect). Patients exhibited significantly higher MSE scores than controls in negative clips of low affective intensity in affective empathy trials. (**C**) Area under the curve (AUC) obtained for negative, low intense clips in patients and controls in affective empathy trials. Positive AUC scores denote more positive subject ratings than target ratings. Compared to controls, patients rated their affect as more positive. Values are means, with standard errors represented by vertical bars, adjusted for BDI-II. * *P* < 0.05 (corrected).

Affective empathy was indexed by the MSE between targets’ and subjects’ self-ratings of emotion. A mixed-design ANCOVA found a significant main effect of Intensity (F(1,41) = 13.18, *P* = 0.001, *η^2^* = 0.24), but not Valence (F(1,41) = 0.57, *P* = 0.455). Thus, deviations between target and subject affect profiles were significantly larger for stimuli of low affective intensity than for stimuli of high affective intensity. There was no significant Valence by Intensity interaction (F(1,41) = 0.52, P = 0.477). No significant main effect of Group emerged (F(1,41) = 2.31, P = 0.137). Neither a Valence by Group interaction (F(1,41) = 2.90, *P* = 0.096), nor an Intensity by Group interaction (F(1,41) = 0.76, *P* = 0.389) reached statistical significance. However, a significant Valence by Intensity by Group interaction emerged (F(1,41) = 8.70, *P* = 0.005, *η^2^* = 0.18). *Post-hoc* analyses revealed that for negative clips of low affective intensity, patients exhibited significantly higher MSE scores than the controls (*P* = 0.024). No significant group effects occurred for other combinations of Valence and Intensity factor levels (*P* ≥ 0.438). Fig. 2B shows the mean MSE scores from affective empathy trials in patients and controls by stimuli valence (positive or negative) and affective intensity (high or low). The complementary analysis of AUC scores in negative clips of low affective intensity revealed positive AUC scores in patients and controls (see Fig. 2C), meaning that subjects’ self-ratings were less negative than those of targets. However, the AUC scores were significantly higher in patients relative to controls (F(1,41) = 4.30, *P* = 0.044 *η^2^* = 0.10). Together, these findings indicate that patients with insular lesions exhibited specific difficulties with affective empathy for negative emotional states of low affective intensity.

### 3.2. Self-reported empathy

For cognitive empathy, a mixed-design ANCOVA revealed a significant main effect of Group (F(1,41) = 4.60, *P* = 0.038, *η^2^* = 0.10), with patients showing lower scores on both the PT and the FS subscale, as compared to controls. Moreover, there was a significant main effect of Subscale (F(1,41) = 20.23, *P* < 0.001, *η^2^* = 0.33), indicating that both groups scored higher on the PT relative to the FS subscale. The Group by Subscale interaction was not significant (F(1,41) = 3.00, *P* = 0.093).

Regarding affective empathy, a mixed-design ANCOVA yielded similar results with a significant main effect of Group (F(1,41) = 5.41, *P* = 0.025, *η^2^* = 0.12). Relative to controls, patients scored significantly lower on both the EC and PD subscales. Additionally, a significant main effect of Subscale emerged (F(1,41) = 36.29, *P* < 0.001, *η^2^* = 0.47), with both groups scoring significantly higher on the EC than on the PD scale. The Group by Subscale interaction did not rise to a significant level (F(1,41) = 0.02, *P* = 0.883). Mean scores are presented in Table 2.

**Table 2:** Self-reported empathic abilities in patients and controls.

|  | **Patients** | **Controls** |
| --- | --- | --- |
|  | *M ± SE* | *M ± SE* |
| **SPF perspective taking<sup>a</sup>** | 13.92 ± 0.52 | 14.32 ± 0.47 |
| **SPF fantasy scale<sup>a</sup>** | 10.57 ± 0.57 | 12.65 ± 0.52 |
| **SPF empathic concern<sup>b</sup>** | 13.78 ± 0.38 | 14.85 ± 0.35 |
| **SPF personal distress<sup>b</sup>** | 9.73 ± 0.58 | 10.94 ± 0.53 |
| **SPF cognitive empathy** | 12.25 ± 0.42* | 13.48 ± 0.38 |
| **SPF affective empathy** | 11.76 ± 0.35* | 12.89 ± 0.32 |

*Note:* SPF = Saarbrücker Persönlichkeitsfragebogen (German adaptation of the Interpersonal Reactivity Index); *M* = mean (adjusted for BDI-II), *SE* = standard error; <sup>a</sup> cognitive empathy scale;
<sup>b</sup> affective empathy scale; \* *P* < 0.05.

Correlation of behavioural performance (MSE) and self-report data (i.e., total score for cognitive resp. affective empathy) yielded no significant association in any of the groups (patients: all *P* ≥ 0.223, controls: all *P* ≥ 0.083).

### 3.3. Lesion laterality

We found no differences between left and right-injured patients on MSE scores obtained for each combination of Valence and Intensity factor levels in cognitive empathy trials (all *P* ≥ 0.294), or affective empathy trials (all *P* ≥ 0.136). These findings are shown in Supplementary Fig. 2. Moreover, lesion groups did not differ on the total scores for cognitive empathy (*P* = 0.963) or affective empathy (*P* = 0.791) from the SPF (not shown).

### 3.4. Contribution of neuropsychological variables

We tested whether performance on neuropsychological tasks predicted patients’ MSE scores for negative stimuli of low affective intensity in the affective empathy trials. Correlational analyses revealed no significant results (all *P* ≥ 0.078).

## 4. Discussion

The present study is the first, to our knowledge, to systematically explore effects of insular damage on affective and cognitive empathic processing as a function of valence and emotional intensity in a modified version of the Empathic Accuracy Task that aimed to approximate real-world social interactions. The patient group did not differ from the healthy control group in their ability to perceive (i.e. continuously track) emotions of others, indicating intact CE. Remarkably, the patient group did also not differ from the healthy control group in their vicarious sharing of emotions, suggesting spared AE, but with one exception: The patient group exhibited significant difficulties to resonate with negative, low-intensity emotions of targets, tending to less negative emotional ratings than the targets rated their own negative affect. This deficit was not related to trait empathy, neuropsychological or clinical parameters. Moreover, we found no effect of lesion laterality on measures of AE or CE.

The insula has been typically related to the affective component of empathy (Banissy et al., 2012; Marsh, 2018; Shamay-Tsoory, 2011; Shamay-Tsoory & Lamm, 2018; Singer et al., 2009; Wicker et al., 2003; Zaki & Ochsner, 2012), with most pronounced links established for negative emotions (Hillis, 2014). Our finding of the selective deficit on AE for negative, low-intensity stimuli following insular damage supports the disproportionate role of the insula in AE. However, it suggests that insular damage may evoke less pronounced dysfunctions on dynamic, context-rich assessments of AE than in emotion recognition tasks employing simple, static stimuli. Notably, there is evidence from studies of patients with neurodegenerative syndromes affecting the insula (e.g., Huntington’s Disease, Frontotemporal Lobar Degeneration) of spared sensitivity towards contextual cues despite insula dysfunction (Kumfor et al., 2017). These patient samples are known to be impaired in negative emotion recognition from faces, but recent evidence indicates that emotion recognition may be spared in ecologically valid tasks with contextual cues (Baez et al., 2015; Kumfor et al., 2017; Pick et al., 2019), as long as the emotions are displayed unambiguously and consistently (Fernandez-Duque et al., 2010; Kumfor et al., 2017). AE has been shown to rely on a complex, widely distributed network (Bernhardt & Singer, 2012; Fan et al., 2011; Shamay-Tsoory & Lamm, 2018), with the insula as an integral node combining internal, sensory, contextual and trait information with information about uncertainty to generate a subjective feeling state (Bernhardt & Singer, 2012; Singer et al., 2009). Thus, one may assume that our patients were able to detect and vicariously share emotional states observed in targets due to the intact interpretation of context cues by residual networks outside the insular lesions. Moreover, the insula is densely interconnected with cortical and subcortical regions that support and complement the insula in its emotional functions (Berntson et al., 2011; Gogolla, 2017; V. Menon, 2015). For example, the amygdala is known to detect, evaluate and initiate emotional arousal (Gläscher & Adolphs, 2003). The anterior cingulate cortex is involved in the representation of the visceral emotional input in terms of a global emotional feeling(Bernhardt & Singer, 2012). The ventromedial prefrontal cortex supports the integration of the interoceptive with exteroceptive information to generate value representation (Gu & FitzGerald, 2014). Together, these structures may compensate the insula damage, as long as the information is clear and easy to detect. However, during ambiguous emotion display (as it might have been during negative videos of low intensity), information necessary to initiate an adequate affect-sharing response cannot be brought together due to the insula damage. Thus, we suggest a threshold effect of the insular damage in online ratings of AE that may depend on the grade of uncertainty in emotionally relevant information. Future studies may attempt to differentiate the impact of contextual social cues and determine the deficit threshold on AE measures in patients with insular damage.

In our study, patients with insular lesions did not exhibit any CE difficulties in the empathy task. This finding suggests no or at least minimal insular involvement in cognitive aspects of empathy. Indeed, previous functional neuroimaging and lesion-based evidence has indicated that CE involves higher order cognitive functions that primarily rely on a set of brain regions typically involved in theory of mind tasks (e.g., prefrontal cortex, superior temporal sulcus, temporo-parietal junction and the temporal poles) (Hillis, 2014; Shamay-Tsoory, 2011; Zaki et al., 2009). Empathic accuracy is considered as an index of CE (Coll et al., 2017) and we adopted the standard empathic accuracy assessment in our study to track CE. Notably, Zaki et al. (2009) show no link between insula activation and empathic accuracy in healthy subjects. However, Fan et al. (2011) found insula activation in both AE and CE tasks in their meta-analysis of functional neuroimaging studies of empathy. Moreover, recent evidence suggests the insula as a converging zone for interoception, emotion and social cognition (Adolfi et al., 2017). Thus, one may assume that the insula may support some subprocesses of CE, but other major nodes in the CE network may compensate the according deficits after insular loss in a dynamic, naturalistic setting.

Our finding of a selective deficit on AE for negative, low-intensity stimuli is counterintuitive to the previous evidence of insular involvement in empathy for pain (Gu et al., 2012; Singer et al., 2009). However, recent models of empathy outline that empathy for pain cannot be equated with the construct of empathy (Marsh, 2018). Empathy is rather a multi-dimensional construct and each dimension may recruit a different neural system (Marsh, 2018; Zaki & Ochsner, 2012). Indeed, empathy for pain tasks and naturalistic, dynamic tasks approximating real-life social interactions may result in different deficit patterns, as shown by some studies in patients with insular dysfunction due to neurodegeneration (for review see Kumfor et al., 2017)). Thus, insular damage appears to affect specific empathy domains in a different manner. Moreover, different kinds of internal states may differently recruit the insula (Adolphs et al., 2003; Barrett et al., 2016; Vicario et al., 2017). While, for example, pain and disgust may particularly depend on the representation of the somatovisceral state coded in the insula, other emotional states may do so less (Gogolla, 2017; Uddin et al., 2017; Vicario et al., 2017). The positive and negative emotional experiences presented by the targets in the current study did not include direct or implied displays of pain (or disgust). Future research is advised to investigate the engagement of the insula in CE and AE measures for videos of events targeting internal states with prominent somatovisceral components.

The patient group scored significantly lower than the healthy control group on both AE and CE scales of the SPF, a self-report questionnaire on empathy functions. This finding points to a deficit to reflect one’s own belief in one’s empathic characteristic and resonates with previous clinical evidence of impaired self-insight following insular dysfunction (Kumfor et al., 2017). Furthermore, task-based AE and CE measures were not correlated with self-report AE and CE measures in patients, or controls. Such dissociations are quite common in empathy literature (Lee et al., 2011; Oliver et al., 2015; Zaki et al., 2008) and suggest that the two measures capture differential aspects of empathy (self-belief vs. actual ability). Our findings outline that self-reported trait empathy diverges from experimental empathy measures and further emphasize that empathy is indeed a multi-faceted psychological construct. Further studies will help us better understand the relationships among distinct aspects of empathy after insular loss. Moreover, our findings indicate that treatment studies or interventions targeting empathy and related behaviours would benefit from a broad multi-method assessment.

Previous functional neuroimaging and lesion-based evidence has suggested differential roles of the left and right insula in empathy (Boucher et al., 2015; Fan et al., 2011; Leigh et al., 2013; Uddin et al., 2017), though the exact nature of affective asymmetries is not clear. We did not observe laterality effects on experimental or self-report measures of AE or CE. This finding resonates with the recent null finding by Chen et al.(2016) in a group of patients with insular gliomas. Given the importance of empathy for successful social functioning (Shamay-Tsoory, 2011), one may assume ipsi- and contralateral compensation of insula-related functions in real-life interactions. However, our findings do not preclude that sub-processes contributing to AE or CE may be processed in a lateralized manner. For example, representation of autonomic information is known to be lateralized in the insula (Craig, 2014), thus determining lateralized affective processing in the human forebrain (Craig, 2014; Duerden et al., 2013). Physiological measures, such as skin conductance, heart rate or blood pressure may be an interesting avenue to address potentially lateralized sub-processes contributing to empathy functions.

Studies with neuropsychiatric samples with insular dysfunction have shown detrimental effects of cognitive impairment and task difficulty on the performance of emotional tasks (Argaud et al., 2018; Gray & Tickle-Degnen, 2010; Suzuki et al., 2006). In our study, we found that performance on neuropsychological tasks did not predict patients’ performance on negative stimuli of low affective intensity in the affective empathy trials. The patient group scored significantly higher on BDI-II scores, therefore we included the BDI-II as a covariate in our analyses. Thus, we assume that cognitive performance may not explain the selective AE deficit in the patient group.

Our study had several noteworthy strengths. First, our findings are based on a patient sample that has been matched for clinical and neuropsychological aspects and that extends the sample sizes of previous lesion studies. Second, we developed and validated a new video set in German language that may provide a useful tool of empathy investigation in future research. The film clips encompassed non-scripted elaborations on personal real-life emotional situations by non-actors, thus (in contrast to previous lesion-based research) approaching the format of typical social interactions. Accounting for the dynamic nature of empathy (Coll et al., 2017; Zaki & Ochsner, 2009), the task required continuous rating of complex emotional stimuli rather than simple recognition or discrimination of specific emotions, a distinction which appears to be relevant in the understanding of social cognition following insular damage (Kumfor et al., 2017). Moreover, our empathy measures were derived from relating subject ratings to target ratings, which is in contrast to previous studies that compared subjects’ summary emotion ratings to pre-defined “correct” ratings. Furthermore, our task allowed assessing AE and CE within the same paradigm, with AE operationalized as affect sharing, while previous lesion-studies assessed AE rather in terms of emotion discrimination abilities or via questionnaires. Finally, we modelled empathy functions according to valence and emotional intensity. This allowed us a more refined insight into the insular involvement in empathy functions.

There are limitations to be noted. Although we used one of the largest patient samples to date, we were not able to investigate associations between lesion characteristics and empathy functions in detail. Moreover, we did not include measures of social functioning, or prosocial motivation and behaviour. However, such measures would help to understand the impact of empathy deficits after insular damage on the adaptation to real-world demands. With regard to the empathy task, one potential limitation is the small number of videos per experimental condition. Moreover, we used the valence scale to rate emotions in oneself and in the target according to previous concepts of the Empathic Accuracy Task. However, future studies may adopt other ratings, e.g. emotional intensity (Mackes et al., 2018), or the intensity of one emotion category. Finally, future studies should examine the specificity of the current findings to insular damage by including a comparison lesion group (e.g. patients with prefrontal lesions in whom primary CE deficits may be expected).

In sum, our findings provide important insights into the profile of social cognition impairment following insular damage. Empathic functions may be widely spared after insular damage in a naturalistic, dynamic setting, potentially due to the intact interpretation of social context cues by residual networks outside the lesion. The particular role of the insular in AE may evolve specifically in situations that bear higher uncertainty, which points to a threshold role of the insular in online ratings of AE.

## Acknowledgements

All authors appreciate the patients’ contribution to this work and would like to thank them and their families for participation in this study.

## Contributions

TS, WM and OH designed the study. CP, WS, NM and CM assessed MRI/ CT scans and recruited the patient group. OH and MF collected the data. OH, MF, MB and JGT analysed the data. OH wrote the first draft of the manuscript.

