## Supplementary Material for "The effects of emotional valence and intensity on cognitive and affective empathy after insula lesions"

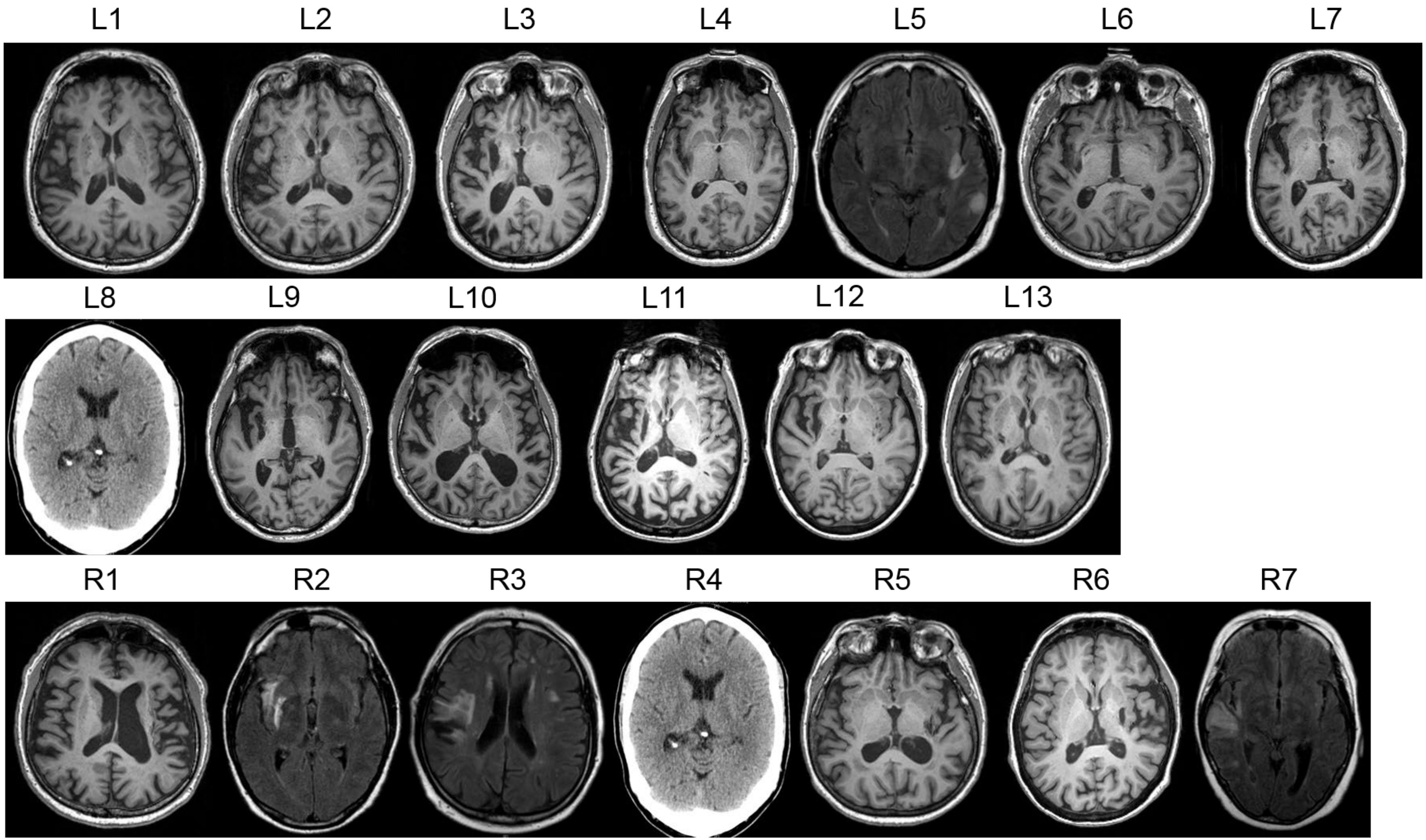


**Supplementary Figure 1:** Overview of individual structural images obtained in left-lesioned (L) and right-lesioned (R) patients. T1-weighted axial MR images were obtained in 14 patients (images in neurological orientation), $T_2$FLAIR MR images in four patients (L5, R2, R3, and R7; images in radiological orientation) and CT images in two patients (L8 and R4; images in radiological orientation).

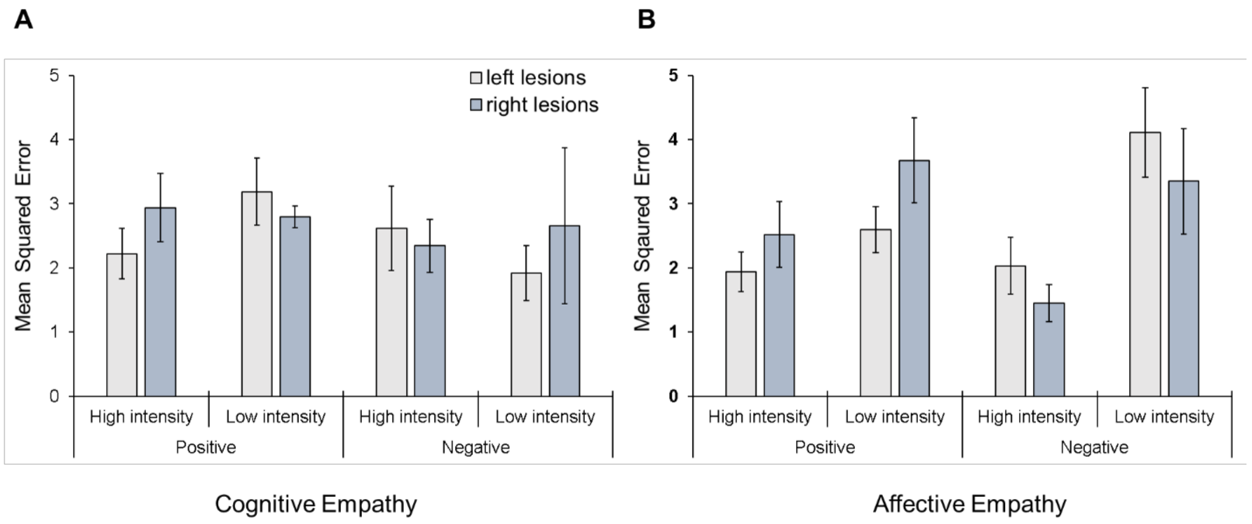


**Supplementary Figure 2:** Mean squared error (MSE) for cognitive empathy trials (**A**) and affective empathy trials (**B**), as a function of lesion group (left-injured, right-injured), valence (positive, negative) and intensity (high, low). No effects of lesion laterality on MSE scores were observed. Values are means, with standard errors represented by vertical bars.

**Supplementary Table 1: Comparative overview of regional lesion volume**

| region | left lesions<br>*M ± SE* | right lesions<br>*M ± SE* | *P* |
|---|---|---|---|
| insula | 1.83 ± 0.37 | 2.57 ± 0.77 | 0.345 |
| basal ganglia | 3.93 ± 1.26 | 3.84 ± 1.19 | 0.961 |
| white matter tracts adjacent to insula and basal ganglia | 2.89 ± 0.88 | 2.35 ± 0.86 | 0.696 |
| thalamus | 0.71 ± 0.38 | 0.60 ± 0.39 | 0.861 |
| limbic structures | 0.14 ± 0.09 | 0.12 ± 0.08 | 0.877 |
| temporal lobe | 4.69 ± 1.28 | 5.99 ± 3.43 | 0.674 |
| central regions | 0.77 ± 0.33 | 1.12 ± 0.96 | 0.676 |
| frontal lobe | 0.49 ± 0.38 | 0.58 ± 0.43 | 0.889 |
| parietal lobe | 1.07 ± 0.78 | 1.19 ± 0.65 | 0.922 |
| occipital lobe | 0.20 ± 0.19 | 0.10 ± 0.10 | 0.465 |
| white matter tracts | 1.89 ± 0.89 | 1.35 ± 0.57 | 0.681 |
| unclassified tissue | 1.60 ± 0.61 | 2.07 ± 0.69 | 0.633 |

*Note:* For each anatomical subdivision, lesion volume is reported in $cm^3$. Standard atlases (AAL atlas, JHU white matter atlas) were used to determine lesion localization and lesion volume on CT and MRI images normalized to MNI space. A claustrum region of interest was added to the analyses, since claustrum is not included in the AAL atlas. Basal ganglia damage covered damage to the caudate nucleus, putamen, pallidum and claustrum. Damage of limbic structures comprised damage to the hippocampus, amygdala, cingulate gyrus and parahippocampal gyrus. Temporal lobe damage referred to damage to the temporal pole, superior, transverse, middle and inferior temporal gyrus. Analyses of damage to central regions, frontal, parietal and occipital lobes followed the anatomical parcellation proposed by Rolls, Joliot, & Tzourio-Mazoyer, 2015. Damage to external and internal capsule, anterior corona radiate, uncinate and superior fronto-occipital fasciculus was subsumed under damage to white matter tracts adjacent to insula and basal ganglia. Damage to all other deep white matter structures included in the JHU white matter atlas was subsumed under damage to white matter tracts. Unclassified tissue refers to damaged regions that were not included in the atlases.

* $P < .05$, uncorrected

Statistical comparisons: *t* tests.

**Supplementary Table 2: Descriptive characteristics of video vignettes used for the investigation of cognitive and affective empathy**

| | | | | | | | Target rating after recording | | | Pilot rating (means) | | | | | |
|---|---|---|---|---|---|---|---|---|---|---|---|---|---|---|---|
| **Condition** | **Valence** | **Intensity** | Vignette Number | Length (sec) | Target gender | Content | Target emotion | Valence [-4; 4] | Intensity [1;9] | Valence [-4;4] | Intensity [1;9] | Clarity of language [1;9] | Comprehensibility of content [1;9] | Authenticity [1;9] | Personable manner [1;9] |
| **Affective empathy** | **positive** | **high** | 1 | 62 | F | holding newborn child | happiness | 4 | 9 | 3.6 | 8.1 | 8.4 | 8.4 | 8.5 | 8.2 |
| | | | 2[a] | 59 | M | family reunion after many years apart after war | happiness | 4 | 9 | 2.7 | 6.3 | 7.8 | 8.1 | 7.5 | 7.4 |
| | | **low** | 3[b] | 61 | F | singing in a caraoke bar | proudness | 3 | 6 | 3.2 | 5.3 | 7.1 | 7.4 | 7.5 | 7.4 |
| | | | 4[c] | 51 | M | funny day at the zoo with own dog | connectedness | 4 | 6 | 3.2 | 5.4 | 7.4 | 8.3 | 7.8 | 8.3 |
| | **negative** | **high** | 5[d] | 68 | F | death of a friend | sadness | -4 | 8 | -1.4 | 7.8 | 8.6 | 8.4 | 8.3 | 7.6 |
| | | | 6 | 50 | M | not recognized by own mother who suffers from dementia | sadness | -4 | 8 | -1.8 | 6.1 | 7.5 | 7.8 | 7.3 | 6.6 |
| | | **low** | 7 | 64 | F | alone on travels | solitude | -4 | 6 | -0.5 | 5.7 | 7.2 | 7.8 | 6.2 | 6.5 |
| | | | 8 | 54 | M | destroying precious teapot on a hotplate | surprise (negative) | -1 | 3 | -1.2 | 3.8 | 5.2 | 6.4 | 7.2 | 6.6 |
| **Cognitive Empathy** | **positive** | **high** | 9[d] | 65 | F | achieve first-time understanding for depression during psychotherapy | relief | 4 | 9 | 1.2 | 8.00 | 8.6 | 8.3 | 8.6 | 7.7 |
| | | | 10[c] | 49 | M | being awarded by a standing ovation as leading artist in a play | proudness | 3 | 9 | 3.1 | 6.7 | 7.7 | 8.1 | 8.0 | 7.8 |
| | | **low** | 11 | 68 | F | discovering a lake on a hot summer day | relief | 2 | 4 | 2.2 | 6.7 | 8.0 | 8.3 | 8.0 | 7.0 |
| | | | 12[a] | 65 | M | spending a day in the allotment garden | relaxation | 2 | 2 | 2.4 | 4.8 | 7.2 | 7.8 | 7.1 | 7.2 |
| | **negative** | **high** | 13 | 58 | F | involuntary separation from the most loved person | solitude | -3 | 9 | -0.80 | 6.5 | 7.1 | 8.0 | 7.0 | 7.4 |
| | | | 14 | 62 | M | deeply hurting a person you love | sadness | -4 | 9 | -1.6 | 7.5 | 8.4 | 8.4 | 7.7 | 7.5 |
| | | **low** | 15[b] | 61 | F | test fear | fear | -1 | 6 | -1.1 | 4.9 | 6.9 | 8.1 | 6.5 | 6.0 |
| | | | 16[a] | 63 | M | best friend distances himself | anger | -3 | 3 | -1.6 | 5.6 | 7.2 | 8.0 | 7.2 | 5.9 |

*Note:* [a-d] indicate video material from respectively the same target; F = female; M = male

### Supplementary Table 3: Discriminative abilities and differences between the affective (AE) and cognitive empathy (CE) video set

| ANOVA | dependent variable | main effect of valence | interaction valence x set |
|---|---|---|---|
| between-subject factors: valence (pos/neg), set (AE/CE) | valence (targets) | F(1,12) = 125.00, ***P* < 0.001** | F(1,12) = 1.80, *P* = 0.205 |
| | intensity (targets) | F(1,12) = 0.03, *P* = 0.857 | F(1,12) = 0.54, *P* = 0.476 |
| within-subject factors: valence (pos/neg), set (AE/CE) | valence (pilot) | F(1,3) = 210.37, ***P* = 0.001** | F(1,3) = 1.59, *P* = 0.297 |
| | intensity (pilot) | F(1,3) = 2.62, *P* = 0.204 | F(1,3) = 0.00, *P* > 0.99 |
| | clarity (pilot) | F(1,3) = 3.62, *P* = 0.153 | F(1,3) = 0.01, *P* = 0.940 |
| | comprehensibility (pilot) | F(1,3) = 1.10, *P* = 0.371 | F(1,3) = 0.55, *P* = 0.512 |
| | authenticity (pilot) | F(1,3) = 6.29, *P* = 0.087 | F(1,3) = 0.33, *P* = 0.604 |
| | personable manner (pilot) | F(1,3) = 13.09, ***P* = 0.036** | F(1,3) = 4.37, *P* = 0.128 |
| **ANOVA** | **dependent variable** | **main effect of intensity** | **interaction intensity x set** |
| between-subject factors: intensity (high/low), set (AE/CE) | valence (targets) | F(1,12) = 0.02, *P* = 0.897 | F(1,12) = 0.02, *P* = 0.897 |
| | intensity (targets) | F(1,12) = 52.45, ***P* < 0.001** | F(1,12) = 2.91, *P* = 0.114 |
| within-subject factors: intensity (high/low), set (AE/CE) | valence (pilot) | F(1,3) = 4.00, *P* = 0.139 | F(1,3) = 0.42, *P* = 0.563 |
| | intensity (pilot) | F(1,3) = 134.66, ***P* = 0.001** | F(1,3) = 0.46, *P* = 0.545 |
| | clarity (pilot) | F(1,3 ) = 12.90**, *P* = 0.037** | F(1,3) = 6.02, *P* = 0.091 |
| | comprehensibility (pilot) | F(1,3) = 5.39, *P* = 0.103 | F(1,3) = 2.37, *P* = 0.221 |
| | authenticity (pilot) | F(1,3) = 7.95, *P* = 0.067 | F(1,3) = 0.03, *P* = 0.878 |
| | personable Manner (pilot) | F(1,3) = 5.10, *P* = 0.109 | F(1,3) = 3.73, *P* = 0.149 |
| **ANOVA** | **dependent variable** | **main effect of sex** | **interaction sex x set** |
| between-subject factors: sex (male/female), set (AE/CE) | valence (targets) | F(1,12) = 0.00, *P* > 0.99 | F(1,12) = 0.283, *P* = 0.604 |
| | intensity (targets) | F(1,12) = 0.54, *P* = 0.476 | F(1,12) = 0.03, *P* = 0.857 |
| within-subject factors: sex (male/female), set (AE/CE) | valence (pilot) | F(1,3) = 0.30, *P* = 0.620 | F(1,3) = 0.96, *P* = 0.399 |
| | intensity (pilot) | F(1,3) = 10.77, ***P* = 0.046** | F(1,3) = 0.74, *P* = 0.454 |
| | clarity (pilot) | F(1,3) = 3.88, *P* = 0.143 | F(1,3) = 0.85, *P* = 0.425 |
| | comprehensibility (pilot) | F(1,3) = 1.29, *P* = 0.339 | F(1,3) = 0.17, *P* = 0.708 |
| | authenticity (pilot) | F(1,3) = 0.08, *P* = 0.791 | F(1,3) = 0.06, *P* = 0.822 |
| | personable Manner (pilot) | F(1,3) = 0.08, *P* = 0.800 | F(1,3) = 0.40, *P* = 0.570 |

*Note:* No statistical differences (*P* < 0.05) were found for both video sets (see right column with interaction results). The sets were found capable to discriminate valence (see main effect of valence for valence ratings) and intensity of emotional content (see main effect of intensity on intensity ratings). Positive videos were rated somewhat higher for personable manner than negative videos ($M_{pos}$ = 7.63, $M_{neg}$ = 6.76). Videos of high affective intensity were rated slightly higher regarding clarity than videos of low affective intensity ($M_{high}$ = 8.01, $M_{low}$ = 7.03). Videos from female targets (F) were rated as somewhat more intense than videos from male (M) targets ($M_F$ = 6.63, $M_M$ = 5.78). Significant findings are printed in bold.
